# BiomiX-Driven Multi-Omics Integration of PRECISESADS Data Reveals Lysophosphatidic Acid and Metabolic Pathway Signatures in B Cells and Immune Macroenvironment in Sjögren’s Disease

**DOI:** 10.64898/2025.11.30.690253

**Authors:** Cristian Iperi, Álvaro Fernández-Ochoa, Nadège Marec, Pierre Pochard, Jacques-Olivier Pers, Guillermo Barturen, Marta Alarcón-Riquelme, PRECISESADS Flow Cytometry Study Group, PRECISESADS Clinical Consortium, Rosa Quirantes-Piné, Isabel Borrás-Linares, Antonio Segura-Carretero, Valérie Devauchelle-Pensec, Divi Cornec, Christophe Jamin, Anne Bordron

## Abstract

**Objective:** Sjögren’s disease (SjD) is a systemic autoimmune disorder characterized by lymphocytic infiltration of exocrine glands, resulting in xerostomia, keratoconjunctivitis sicca, fatigue, arthralgia, and systemic organ involvement. This study aimed to characterize the metabolic and immune dysregulation of SjD using a multi-omics approach, focusing on the metabolic environment and B-cell transcriptomic responses.

**Methods:** Transcriptomic, methylomic, and metabolomic datasets from whole blood, plasma, and urine of 293 SjD patients and 508 controls were analyzed from the PRECISESADS study. B-cell transcriptomes were included to link systemic metabolic alterations to cell-intrinsic immune programs. Multi-omics factor analysis (MOFA) was used to integrate data and identify discriminant molecular drivers.

**Results:** Multi-omics integration revealed metabolic rewiring involving the urea cycle, glutamine/arginine metabolism, and NAD⁺ depletion linked to interferon signaling. Among the strongest contributors, plasma lysophosphatidic acids (LPA) emerged as key discriminants associated with interferon-driven activation. B-cell transcriptomes showed upregulation of LPA-related genes (CERS6, INPP1, TRIP6), and its receptor LPAR6. Importantly, in this study LPAR6 protein expression was confirmed in B cells for the first time. Secondary findings included alterations in sphingosine-1-phosphate (S1P) metabolism, suggesting a broader lysophospholipid signaling axis.

**Conclusions:** This study identifies the LPA–LPAR6 signaling axis as a potential metabolic driver of B-cell activation and interferon-associated inflammation in SjD, highlighting a previously unrecognized immunometabolic pathway. These findings highlight LPA–LPAR6 as a candidate target for therapeutic modulation in SjD, while also implicating S1P signaling as a complementary regulatory mechanism.

## 1. Introduction

Sjögren’s disease (SjD) is a chronic systemic autoimmune disorder affecting 1 in 1,664 individuals, predominantly women (16:1), with a mean onset at 56 years[1]. Beyond xerostomia and xerophthalmia, SjD presents with pulmonary, renal, neurological, and arthritic involvement[2,3] and carries a markedly increased risk of non-Hodgkin lymphoma, 10.5 to 48-fold higher than the general population, reaching 15.4% in men[4]. These features highlight the need for improved mechanistic insights and therapies. Central to disease pathogenesis is B-cell hyperactivation: B cells form ectopic germinal center–like structures, produce autoantibodies (anti-SSA/Ro, anti-SSB/La), and drive inflammation[5]. The relevance of B cells is supported by B-cell–directed therapies, including the phase III agent ianalumab[6], though heterogeneous responses indicate unresolved mechanisms of activation.

Immunometabolism has emerged as a major regulator of immune function. Activated B cells rely on glycolysis, glutaminolysis, and mitochondrial respiration, and in SjD display increased glucose use and OXPHOS; inhibiting glycolysis, mTORC1, or respiration reduces activation and plasma cell differentiation[7]. Systemic metabolites may also modulate immune thresholds[8], underscoring the need to integrate cellular and systemic metabolic cues.

High-throughput multi-omics approaches enable discovery of shared molecular drivers across regulatory layers[9,10]. In SjD, multi-omics clustering has already revealed biologically distinct patient groups[11]. Building on this, the present study applies Multi-Omics Factor Analysis (MOFA) [12], previously applied in SLE[13], to integrate transcriptomic, methylomic, and metabolomic datasets. By combining plasma/urine metabolomics with whole-blood, PBMC, and B-cell transcriptomics, the analysis links systemic metabolic signals to cell-intrinsic B-cell pathways. The objective of this study is to define metabolic and transcriptional programs underlying SjD pathogenesis with a particular focus on B-cell biology, and identify mechanisms that may lead future therapeutic strategies.

## 2. Material and methods

### 2.1 Samples and cohort selection

The cross-sectional PRECISESADS cohort (NCT02890121) was designed to reclassify systemic autoimmune diseases (SAD) using clinical and multi-omics data. Patient selection and quality control procedures were previously described[14]. The study complied with ICH-GCP and the Declaration of Helsinki (2013). All participants provided informed consent, and the protocol was approved by ethics committees of the 19 participating institutions. This substudy focused on SjD, defined by the 2002 American-European Consensus Group criteria[15]. Whole blood samples from SjD (n = 293) and healthy controls (CTRL, n = 508) were split into two aliquots. The first contained all immune cell types; the second was used for additional analyses. Purified B cells were isolated from part of the second aliquot (41 SjD, 27 CTRL). Plasma and urine were collected from 99 paired samples (45 SjD, 54 CTRL) for metabolomics. Whole blood fractions were used for methylomics (282 SjD, 364 CTRL). Metadata for whole blood are summarized in **Table 1**.

**Table 1.**
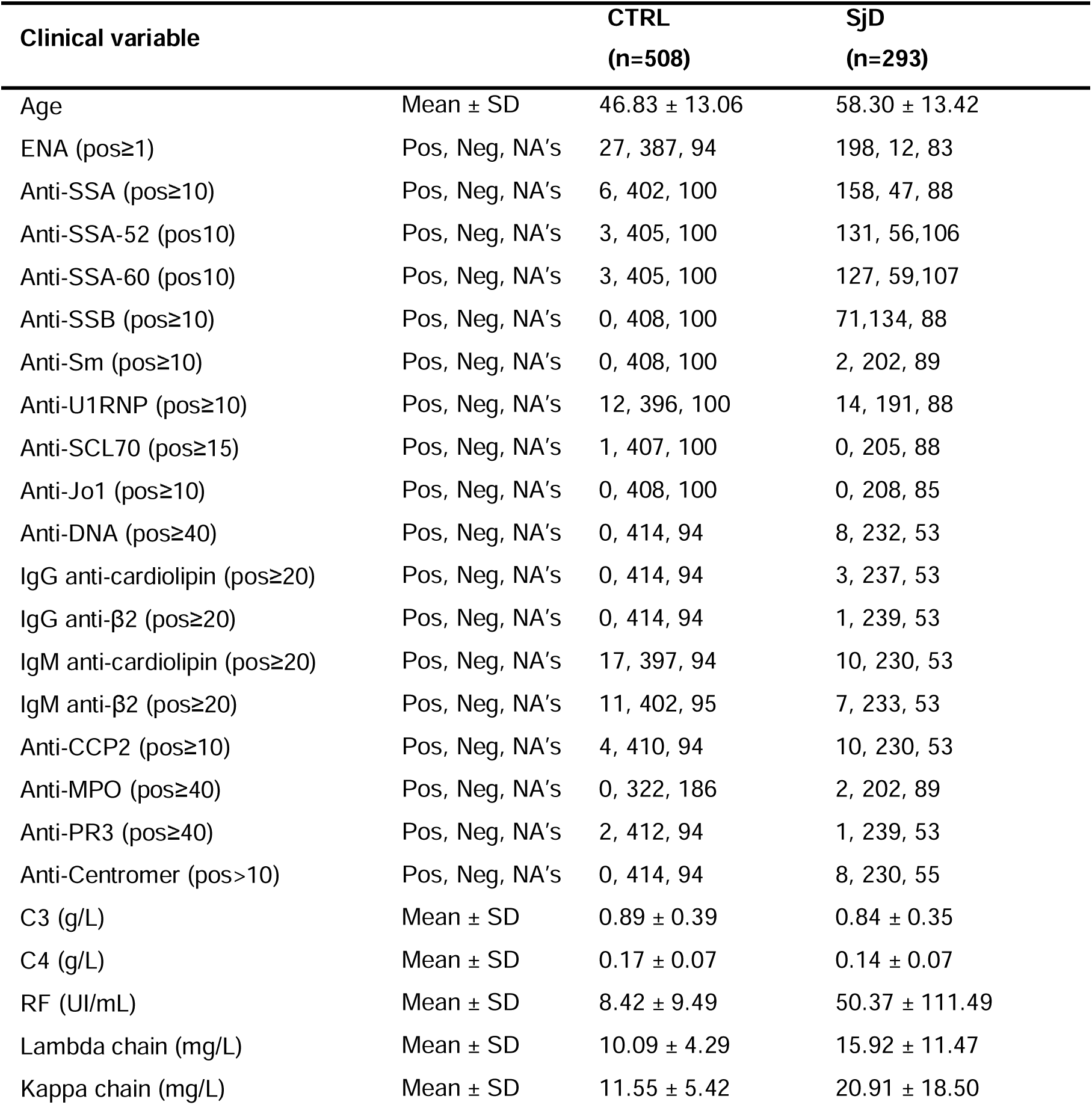
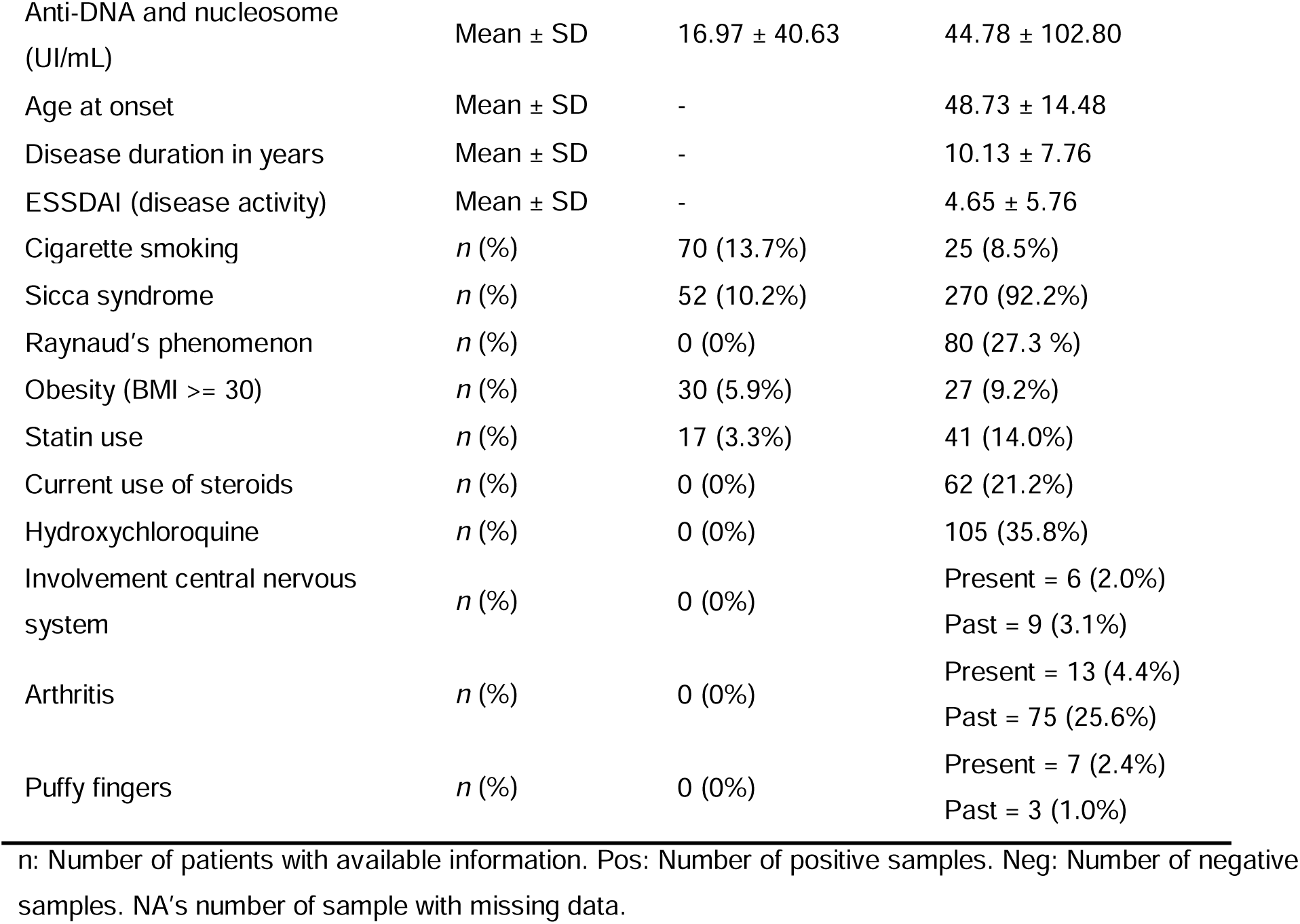
Healthy controls (CTRL) and Sjögren’s disease (SjD) patient characteristics.

| Clinical variable |  | CTRL<br>(n=508) | SjD<br>(n=293) |
| --- | --- | --- | --- |
| Age | Mean ± SD | 46.83 ± 13.06 | 58.30 ± 13.42 |
| ENA (pos≥1) | Pos, Neg, NA's | 27, 387, 94 | 198, 12, 83 |
| Anti-SSA (pos≥10) | Pos, Neg, NA's | 6, 402, 100 | 158, 47, 88 |
| Anti-SSA-52 (pos10) | Pos, Neg, NA's | 3, 405, 100 | 131, 56, 106 |
| Anti-SSA-60 (pos10) | Pos, Neg, NA's | 3, 405, 100 | 127, 59, 107 |
| Anti-SSB (pos≥10) | Pos, Neg, NA's | 0, 408, 100 | 71, 134, 88 |
| Anti-Sm (pos≥10) | Pos, Neg, NA's | 0, 408, 100 | 2, 202, 89 |
| Anti-U1RNP (pos≥10) | Pos, Neg, NA's | 12, 396, 100 | 14, 191, 88 |
| Anti-SCL70 (pos≥15) | Pos, Neg, NA's | 1, 407, 100 | 0, 205, 88 |
| Anti-Jo1 (pos≥10) | Pos, Neg, NA's | 0, 408, 100 | 0, 208, 85 |
| Anti-DNA (pos≥40) | Pos, Neg, NA's | 0, 414, 94 | 8, 232, 53 |
| IgG anti-cardiolipin (pos≥20) | Pos, Neg, NA's | 0, 414, 94 | 3, 237, 53 |
| IgG anti-β2 (pos≥20) | Pos, Neg, NA's | 0, 414, 94 | 1, 239, 53 |
| IgM anti-cardiolipin (pos≥20) | Pos, Neg, NA's | 17, 397, 94 | 10, 230, 53 |
| IgM anti-β2 (pos≥20) | Pos, Neg, NA's | 11, 402, 95 | 7, 233, 53 |
| Anti-CCP2 (pos≥10) | Pos, Neg, NA's | 4, 410, 94 | 10, 230, 53 |
| Anti-MPO (pos≥40) | Pos, Neg, NA's | 0, 322, 186 | 2, 202, 89 |
| Anti-PR3 (pos≥40) | Pos, Neg, NA's | 2, 412, 94 | 1, 239, 53 |
| Anti-Centromer (pos>10) | Pos, Neg, NA's | 0, 414, 94 | 8, 230, 55 |
| C3 (g/L) | Mean ± SD | 0.89 ± 0.39 | 0.84 ± 0.35 |
| C4 (g/L) | Mean ± SD | 0.17 ± 0.07 | 0.14 ± 0.07 |
| RF (UI/mL) | Mean ± SD | 8.42 ± 9.49 | 50.37 ± 111.49 |
| Lambda chain (mg/L) | Mean ± SD | 10.09 ± 4.29 | 15.92 ± 11.47 |
| Kappa chain (mg/L) | Mean ± SD | 11.55 ± 5.42 | 20.91 ± 18.50 |
| Anti-DNA and nucleosome (UI/mL) | Mean ± SD | 16.97 ± 40.63 | 44.78 ± 102.80 |
| Age at onset | Mean ± SD | - | 48.73 ± 14.48 |
| Disease duration in years | Mean ± SD | - | 10.13 ± 7.76 |
| ESSDAI (disease activity) | Mean ± SD | - | 4.65 ± 5.76 |
| Cigarette smoking | *n* (%) | 70 (13.7%) | 25 (8.5%) |
| Sicca syndrome | *n* (%) | 52 (10.2%) | 270 (92.2%) |
| Raynaud's phenomenon | *n* (%) | 0 (0%) | 80 (27.3 %) |
| Obesity (BMI ≥ 30) | *n* (%) | 30 (5.9%) | 27 (9.2%) |
| Statin use | *n* (%) | 17 (3.3%) | 41 (14.0%) |
| Current use of steroids | *n* (%) | 0 (0%) | 62 (21.2%) |
| Hydroxychloroquine | *n* (%) | 0 (0%) | 105 (35.8%) |
| Involvement central nervous system | *n* (%) | 0 (0%) | Present = 6 (2.0%)<br>Past = 9 (3.1%) |
| Arthritis | *n* (%) | 0 (0%) | Present = 13 (4.4%)<br>Past = 75 (25.6%) |
| Puffy fingers | *n* (%) | 0 (0%) | Present = 7 (2.4%)<br>Past = 3 (1.0%) |

*n*: Number of patients with available information. Pos: Number of positive samples. Neg: Number of negative samples. NA's number of sample with missing data.

### 2.2 Transcriptome data generation

The whole blood and sorted B cell samples were sequenced using a Novaseq 5000 and a NextSeq 500, respectively, with an average coverage per sample of 11.4 Million reads for the former and 29.9 Million reads for the latter. The FastQ files were aligned to the UCSC *Homo sapiens* reference genome (hg19) and annotated to GENCODE 19 with STAR v2.5.2[16] using two-pass mapping strategy with default parameters. Gene quantification was performed using RSEM v1.2.31[17].

### 2.3 Metabolomics data generation

Untargeted metabolomics followed previously published PRECISESADS protocols[18]. HPLC-ESI-QTOF-MS analysis used a reverse-phase C18 column (Zorbax Eclipse Plus, 2.1 mm×150 mm, 3.5 μm) at 25 °C on a 6540 UHD Accurate-Mass Q-TOF (Agilent) equipped with a Jet Stream dual ESI interface operated in positive-ion mode. Data were converted to .mzML with ProteoWizard MSConvert(v3.0.)[19]. Parameter optimization used IPO(v1.10.0)[20]; peak picking, alignment, and grouping were performed with XCMS (v3.6.0)[21], batch correction by batchCorr v0.2.1(v0.2.1)[22], and feature grouping by RamClustR[23]. QC pools prepared by mixing equal aliquots of the samples and injected every 6 experimental samples monitored instrument performance, reproducibility, drift, and low-quality features (RSD < 30%); blanks were used to remove contaminants. After processing, 531 (plasma) and 1,022 (urine) molecular features were retained for comparison.

### 2.5 Methylomics data generation

As described in PRECISESADS study[11]. DNA was extracted using a magnetic-bead protocol (Chemagic DNA Blood Kit; PerkinElmer) from K2EDTA tubes (lavender cap, BD Vacutainer) on 3□ml. Two micrograms of DNA were hybridized to the Infinium HumanMethylation450K array (Illumina) covering more than400,000 CpGs. Data were analyzed in BiomiX with thresholds of |Δβ| ≥ 0.1 and adjusted p ≤ 0.05.

### 2.6 MOFA analysis

MOFA was run in BiomiX[24] with automated factor optimization. Tuning halted when ≥3 models showed <1% variance explained for the last factor. The statistical discrimination between the two groups is determined for each calculated factor in each model and only the top three models discriminating are maintained. Mann–Whitney tests with FDR correction assessed group separation by factor values[25]. The 20-factor model of MOFA had the highest number of discriminant factors (factors 4, 8, and 10) and was selected for the downstream analysis.

### 2.7 Clinical data analysis and omics data batch effect check

Sixty clinical variables and 19 autoantibodies were correlated with significant MOFA factors using Pearson tests, while 160 binary clinical features were assessed with Wilcoxon tests. Nominal p-values were adjusted with the Benjamini–Hochberg method. Age, gender, corticosteroid use, and hydroxychloroquine treatment were evaluated across transcriptomic, metabolomic, and methylomic datasets by testing the first four principal components (t-tests for categorical variables; ANOVA for age groups <35, 35–50, >50). No Bonferroni-corrected p-values indicated detectable effects.”

### 2.8 BiomiX parameters

#### Transcriptomics-parameters

BiomiX (GitHub: IxI-97/BiomiX) processed transcriptomic, methylomic, and metabolomic data and integrated them into MOFA. Differential gene expression (DGE) used thresholds of |log₂FC| ≥ 0.5 and adjusted p ≤ 0.05. Gene set enrichment was assessed with GSEA v4.40 using Reactome and Gene Ontology databases. IFN-α status was assigned using a 26-gene panel [SIGLEC1, IFIT3, IFI6, LY6E, MX1, USP18, OAS3, IFI44L, OAS2, IFIT1, EPSTI1, ISG15, RSAD2, HERC5, OAS1, IFI44, SPATS2L, PLSCR1, IFI27, RTP4, EIF2AK2, GBP1, IRF1, SERPING1, CXCL10, and FCGR1A[13,26], defining positivity when >10 genes had Z>2 or >20 genes had Z>1. Clustering parameters matched heatmap settings (Euclidean distance, Ward.D). For DGE, only B-cell samples with digital purity >90% and IFN-α–negative CTRLs were included (28 SjD, 19 CTRL; **Table S1**).

#### Metabolomics-parameters

Significant features met |log₂FC| ≥ 0.5 and adjusted p ≤ 0.05. MS¹ annotation used 15-ppm tolerance, positive-ion mode, and common adducts ([M+H]⁺, [M+2H]²⁺, [M+Na]⁺, [M+K]⁺, [M+NH₄]⁺, [M+H−H₂O]⁺), with HMDB, LipidMaps, Metlin, and KEGG queried via CEU Mass Mediator. MS² annotation by fragmentation spectra (15-ppm tolerance; .mzML files) used HMDB, MoNA, and MassBank. Significant m/z values are listed in **Table S2**.

#### MOFA-parameters

MOFA was run in fast mode (5,000 iterations) with a loading threshold of 0.5. The optimal model was selected automatically based on the number of discriminant factors distinguishing SjD from CTRLs; only samples with all three omics layers were used (**Table S3**). Pathway enrichment was applied to top contributors, retaining the 10 most significant pathways (adjusted p<0.05) per factor. Associations with numerical and binary clinical variables were tested, and biological interpretation was supported by screening the top 300 PubMed abstracts per discriminant factor.

### 2.8 LPAR6 protein flow cytometry validation

B cells were isolated from apheresis residues (EFS Rennes) by CD19-positive magnetic selection (Miltenyi, ref:130-117-034). Purity was confirmed by flow cytometry. For LPAR6 staining, cells were incubated with rabbit anti-LPAR6 antibody (Alomone; ref:ALR-036) or isotype control for 30 min at 4 °C, then stained with donkey anti-rabbit IgG Alexa Fluor 488 (BioLegend) and anti-CD19 APC-AF700 (Beckman Coulter) for 15 min at 4 °C. Acquisition was performed on a BD FACSymphony A3 and analyzed with FlowJo. Kruskal–Wallis tests (R) compared stained, isotype, and secondary-only conditions.

## 3. Results

### 3.1 The Fluidic Macroenvironment: Plasma and Urine Metabolomics

The disease macroenvironment of SjD can be subdivided into a cellular macroenvironment, represented by the whole-blood transcriptome, and a fluidic macroenvironment, comprising plasma and urine metabolomes. To initiate the characterization of SjD patients, the plasma and urine metabolites were profiled first to define the metabolic environment to which PBMCs, and in particular B cells, are exposed.

Plasma metabolomics data revealed an increase in citrulline, a metabolite involved in mediating inflammation in autoimmunity[27] (**Figure 1**). Lipid metabolism was prominently perturbed, with increased levels of sphingosine-1-phosphate (S1P), as well as LysoPC(22:6) and LysoPC(22:5), the choline-containing direct precursors of lysophosphatidic acids (LPA), together with cholesterol glucuronide. Several carnitines were altered, with hexanoyl- and octanoyl-L-carnitine being increased, whereas linoelaidyl-carnitine was reduced. Consistently, trimethyl-lysine, the initial metabolite in the carnitine biosynthesis pathway, was also elevated in plasma. Urine metabolomics data revealed a reduction in methylxanthine, a caffeine-derived metabolite, and an increase in phenylacetylglutamine and 2-phenylglycine. Trimethyl-lysine and sphingosine were also detected in urine, but both were reduced in SjD patients compared to CTRLs. Importantly, the plasma lipidomic profile revealed two distinct peaks suggesting a potential increase in LPA. LPA is a bioactive lipid mediator known to drive inflammation and, in cancer, has been reported to promote immune evasion and migration[28]. Its involvement in autoimmunity has been described in SLE and, more recently, in systemic sclerosis (SSc)[13,29]. In SSc, two phase 3 clinical trials (NCT06003426 and NCT06025578) are evaluating the LPA receptor 1 inhibitor BMS-98627 for idiopathic pulmonary fibrosis (IPF) and interstitial lung disease (ILD), raising the question of whether elevated LPA may play a similar role in the immune dysregulation observed in SjD. To determine whether the transcriptional programs of circulating immune cells mirror this pro-inflammatory, lipid-enriched environment, the cellular macroenvironment was examined through whole-blood transcriptomic and methylomic profiling.

**Figure 1.**
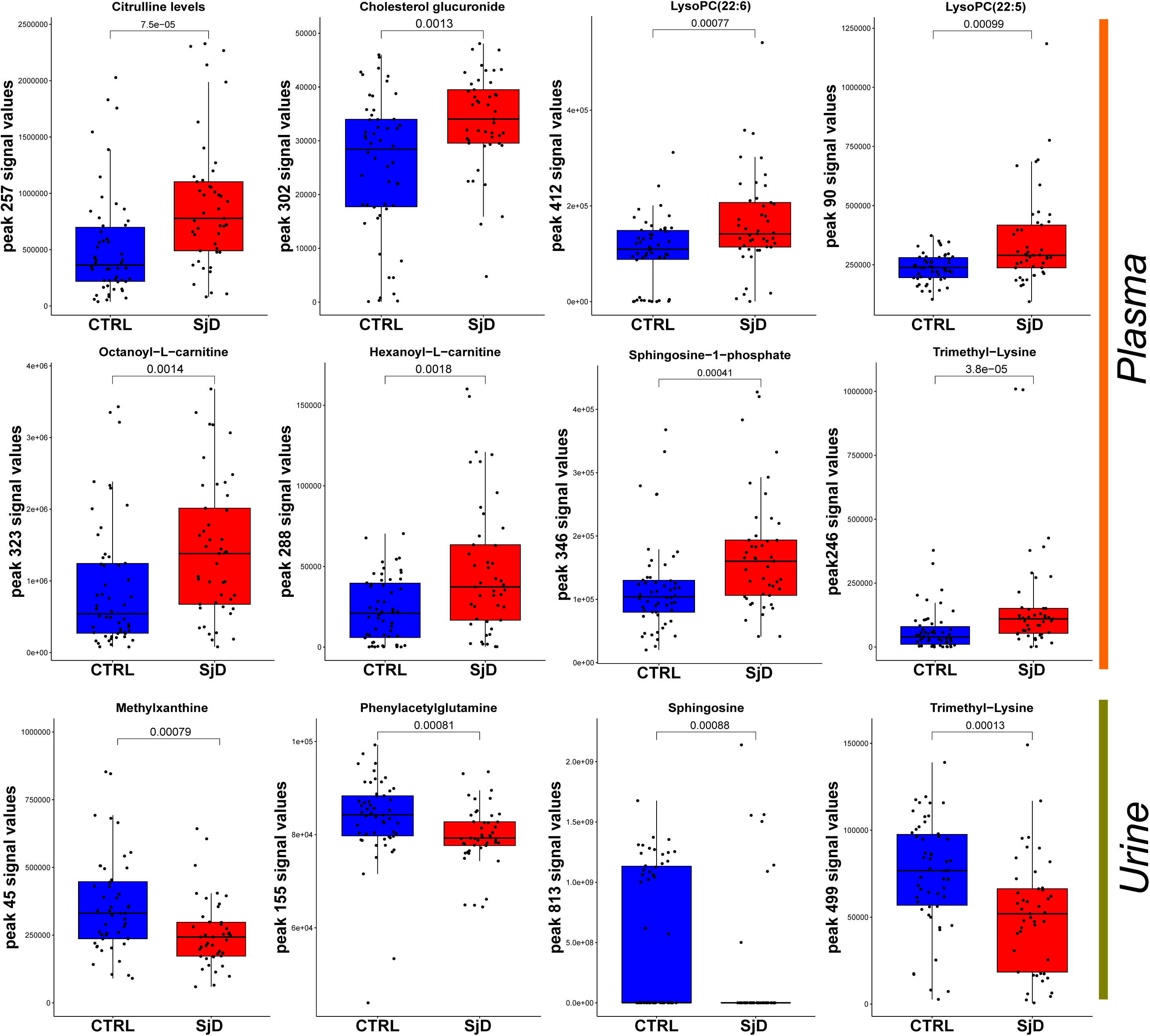
Statistically significant metabolites altered in SjD Plasma and Urine. Levels of metabolites in plasma and urine of control (CTRL) Sjôgren’s patients (SjD) are depicted in blue and red, respectively. Significance was calculated using the Mann-Whitney test

### 3.2 The Cellular Macroenvironment: Whole-Blood Transcriptomics and Methylomics

Whole-blood transcriptome analysis identified 1,074 overexpressed and 422 underexpressed genes in SjD patients compared with CTRLs (**Figure 2A–B**). Among the most strongly upregulated transcripts were interferon-stimulated genes (ISGs), including IFI44L, RSAD2, and OAS3. GSEA confirmed this pattern, revealing activation of type I (α, β) and type II (γ) interferon pathways, together with cytokine signaling via Toll-like receptors (TLRs), macrophage activation, and antigen presentation through MHC class I. GSEA also highlighted signatures of cellular activation and stress, such as oxidative and endoplasmic reticulum stress, known to be associated with apoptosis and autophagosome formation, respectively.

**Figure 2.**
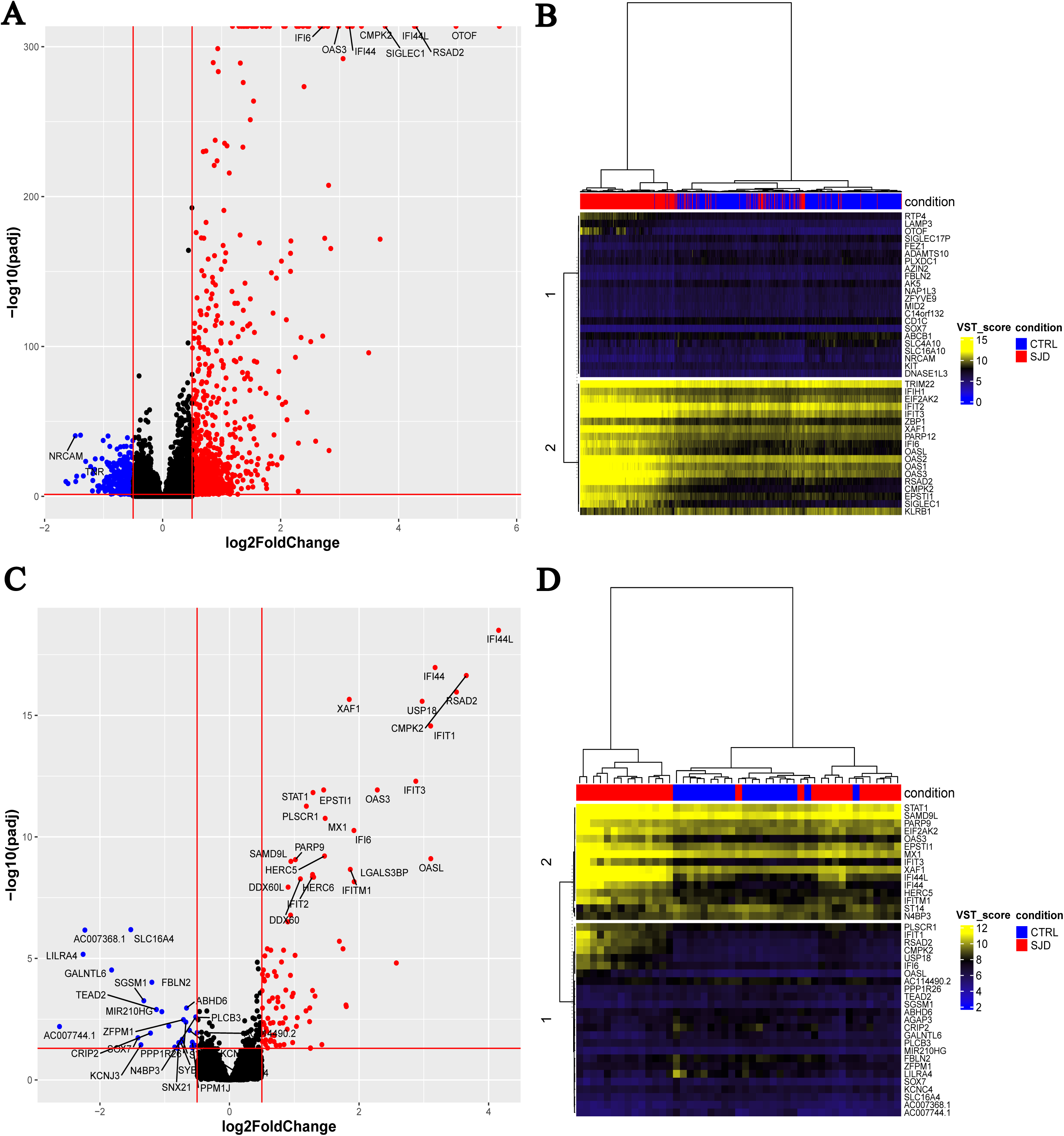
Transcriptomic alterations in whole blood and B cells of SjD patients compared with controls (CTRL) in the PRECISESADS dataset. **A)** Volcano plot of differential gene expression in whole blood. Downregulated and upregulated genes are shown in blue and red, respectively. The top 25 upregulated and top 2 downregulated genes are highlighted. Red lines indicate the thresholds of significance: |Log2FC| > 0.5 and adjusted *p* < 0.05. **B**) Heatmap of the top 20 downregulated and top 20 upregulated genes in whole blood. Blue and red squares above the heatmap indicate CTRL and SjD patients, respectively. Distances were calculated using the Euclidean method, with complete linkage clustering. Gene expression values were normalized using the variance stabilizing transformation (VST). **C**) Volcano plot of DGE in B cells, using the same parameters as in (A). **D**) Heatmap of the top 20 downregulated and top 20 upregulated genes in B cells, generated as in (B).

Beyond immune activation, the expression patterns suggested a marked metabolic activation. Genes involved in glucose and amino acid transport and utilization, particularly for arginine and glutamine, were overexpressed. The ARG1 and ASS1 urea cycle enzymes required for citrulline production were overexpressed in line with the increased citrulline observed in plasma. Similarly, genes encoding transporters for sterols and long-chain fatty acids were upregulated, together with those driving biosynthetic pathways for glycolipids, phosphatidylcholine, and ceramide. Genes involved in both degradation and biosynthesis of NAD+, as well as processes related to cell adhesion and integrin signaling, were also upregulated, supporting the presence of a metabolically and functionally active cellular state. All GSEA and differentially expressed genes are reported in **Supplementary file 1**.

Methylomic analysis revealed 15 demethylated CpG sites, most of which were associated with ISGs such as MX1 (three sites), IFI44L, NLRC5, IFIT1, PARP9 (two sites), PLSCR1, IFITM1, and CMPK2, in addition to three sites within HLA-A and one within HLA-C (**Figure S1**). These epigenetic changes were concordant with the transcriptomic results, as reduced methylation of these genes is consistent with their increased expression. Together, these results reveal a cellular environment characterized by interferon-driven activation and metabolic rewiring. The analyses then focused on B lymphocytes to determine whether these transcriptional and metabolic changes were reflected at the level of this key cell type.

### 3.3 B Cell Transcriptomics: ISG and Lysophosphatidic Pathway Activation

After characterizing the overall SjD environment, including changes in whole blood cells and circulating metabolites, the B lymphocyte-specific transcriptome was analyzed to identify both unique and shared alterations relative to the PBMC profile. B cell transcriptomic analysis revealed 92 overexpressed and 29 underexpressed genes (**Figure 2C–D**). Although GSEA analysis was inconclusive (**Supplementary file 1**), pathway analysis using EnrichR confirmed activation of interferon-mediated signaling pathways (GO:0140888, GO:0035456, GO:0035455), cytokine signaling (GO:0019221), and regulation of B cell proliferation (GO:0030888).

Among the overexpressed genes were those associated with nucleotide depletion (e.g., CMPK2, RSAD2) and several schafen proteins (SLFN5, SLFN11, SLFN12L, SLFN13). Notably, LPAR6, encoding a receptor for LPA, was also upregulated with other genes related to LPA metabolism and signaling (APOL6, INPP1, CERS6, TRIP6). This finding suggested that B cells may be primed to respond to the elevated LPA levels identified in plasma metabolomics, potentially linking the inflammatory lipid environment to B cell activation. To explore whether these changes represented a shared driver across multiple molecular layers, a multi-omics integration analysis was performed.

### 3.4 Multi-omics integration in Sjögren’s data reveals LPA-Interferon axis

To improve the understanding of common processes underlying SjD, the multi-omics factor analysis (MOFA) was applied, which enables joint analysis of transcriptomic, methylomic, and metabolomic data to identify shared sources of variation. The optimal MOFA model was selected by iteratively increasing the number of factors until additional factors explained <1% of novel variance. The 20-factor model was chosen, containing three factors (4, 8, and 10) that significantly discriminated SjD from CTRLs and showed the highest discriminatory power (**Figure 3A–C**). Factors 4 and 10 were increased in SjD patients, whereas factor 8 was decreased. Factor 4 was primarily driven by whole-blood transcriptomics (13.89% variance explained) and methylomics (2.23%), with a minor contribution from B cell transcriptomics (0.26%). Factor 8 was mainly explained by whole-blood transcriptomics (6.72%), B-cell transcriptomics (0.43%), and plasma metabolomics (0.17%), while factor 10 captured variance largely from whole-blood (3.08%) and B-cell transcriptomics (1.92%). Variance explained for all 20 factors is reported in **Table S4**, and contributors by omics type are in **Table 4**. The anti-SSB, anti-SSA52, anti-SSA60, anti-ENA autoantibodies were correlated with SjD condition in factors 8 and 10, with anti-JO1, anti-U1RNP, IgG anti-cardiolipin (CLG) exclusive for factor 8 and Rheumatoid factor (RF) in factor 10. Factor 4 correlated with SjD condition in SSA, ENA, PR3 and SCL70 (**Supplementary File 1**).

**Figure 3.**
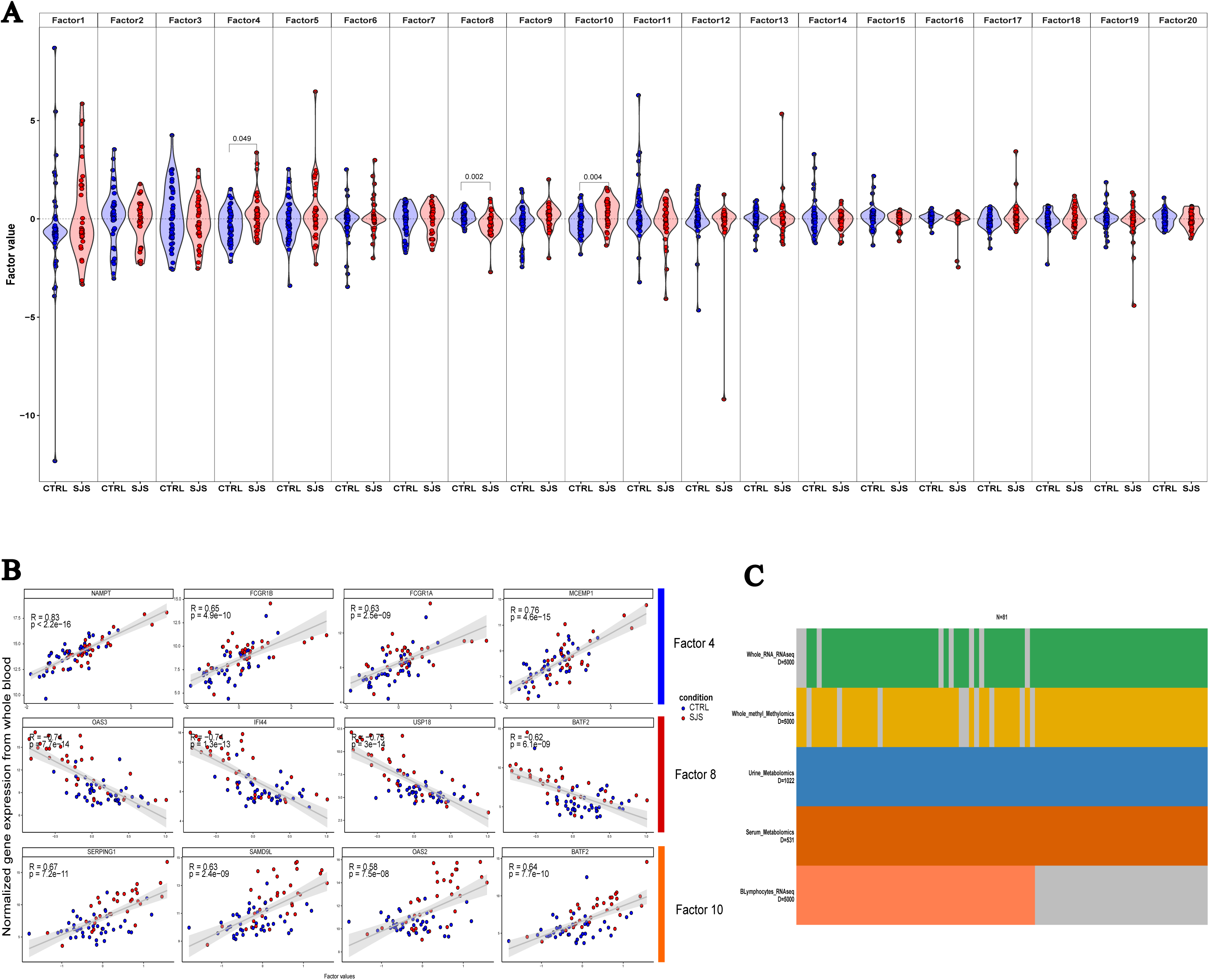
Identification of variation among multi-omics data from Sjögren’s disease patients using Multi-Omics Factor Analysis (MOFA) factors. **A)** Violin plot representing SjD (red) and control sample (CTRL; blue) distribution based on each MOFA factor’s value. **B)** Scatterplot of the top whole-blood 4 genes contributing to Factor 4, 8 and 10. **C)** Representation of the MOFA datasets integrated into the analysis. The numbers of features (N) used in the MOFA integration include peak signals from urine metabolomics and plasma metabolomics, genes from whole-blood transcriptomics and B-cell transcriptomics, and CpG methylation from whole-blood methylomics among a total of 81 samples.

**Table 4.**
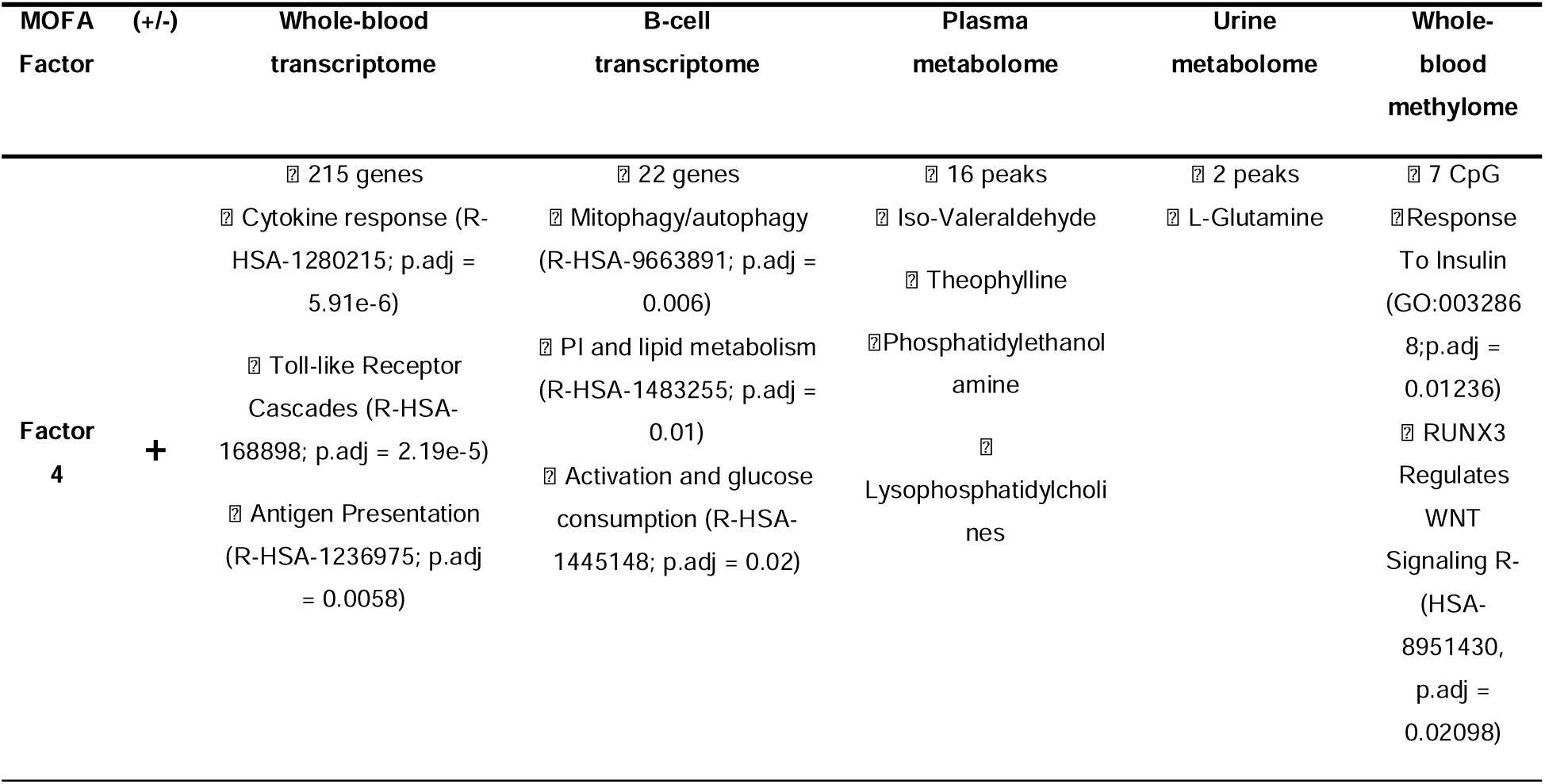
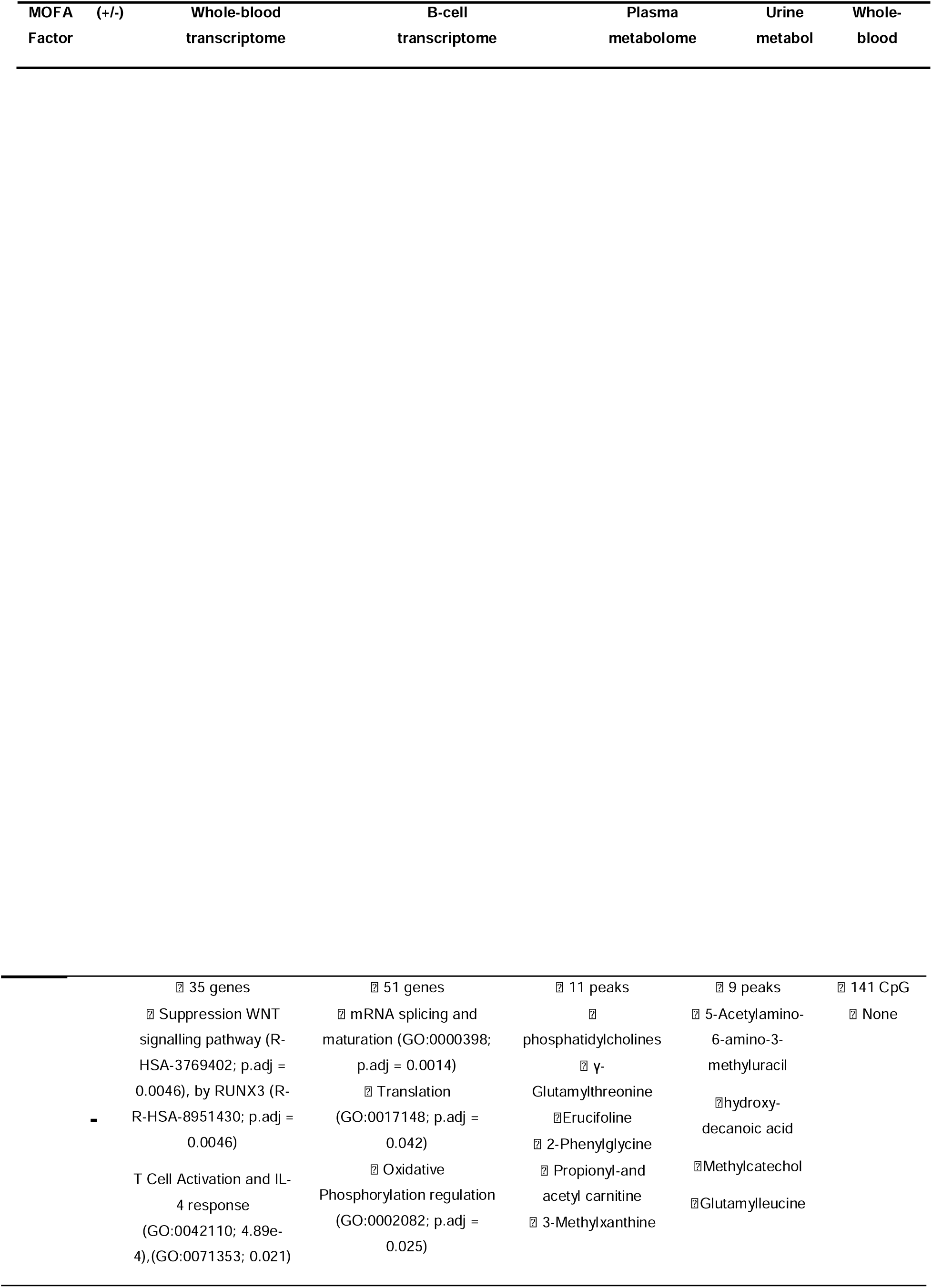
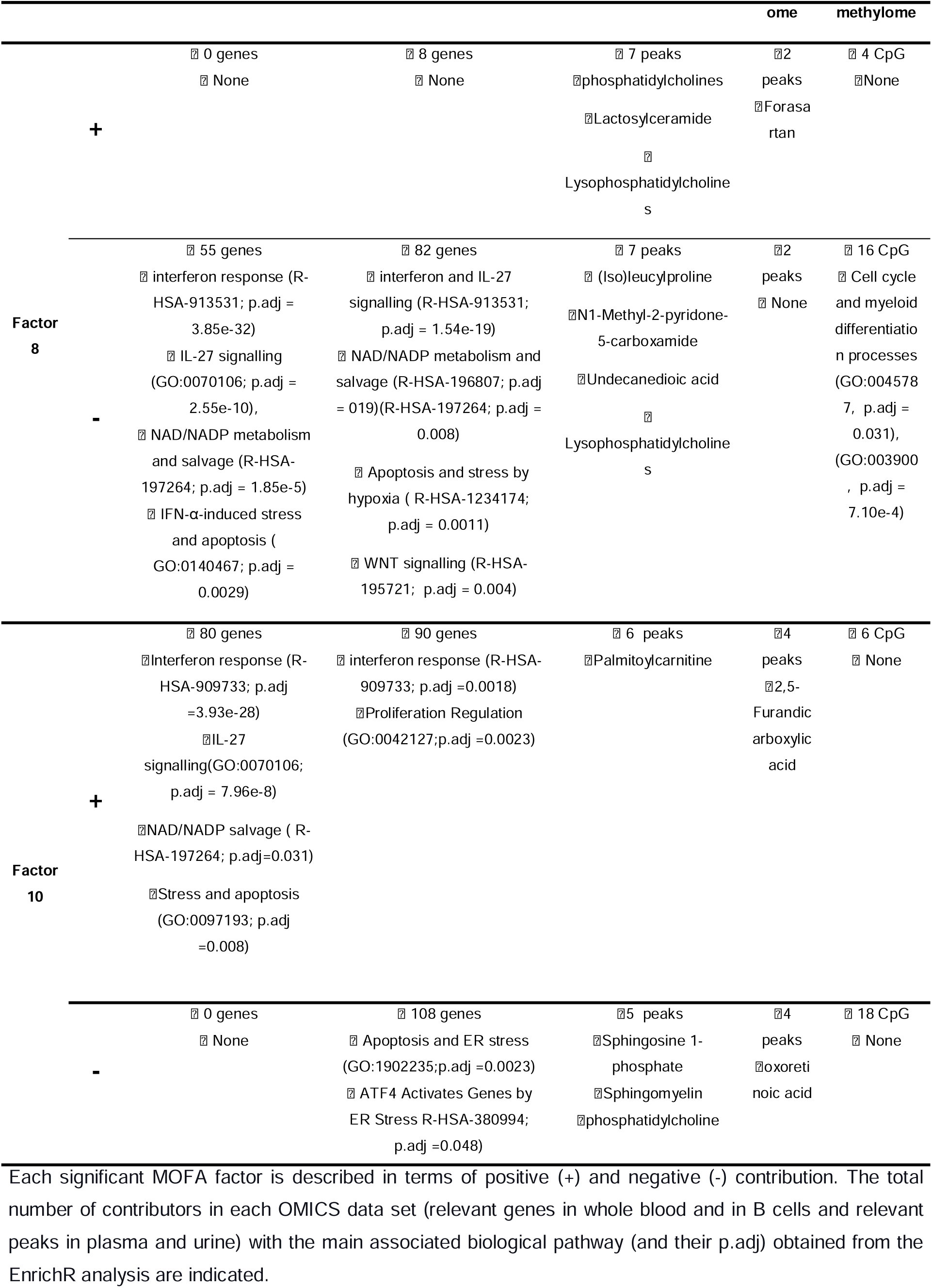
Biological processes of significant MOFA factors and significant contributors.

EnrichR pathway enrichment analysis on top contributors revealed that factor 4 was associated with cellular activation, including Toll-like receptor cascades (R-HSA-168898), neutrophil degranulation (R-HSA-6798695), and inflammatory responses in whole blood, mirrored in B cells with stress-associated lipid synthesis (R-HSA-1483255), mitophagy (R-HSA-5205647), and apoptosis (GO:0042981). Negative contributors included mRNA splicing and processing (GO:0000398, GO:0006397), oxidative phosphorylation regulation (GO:0002082), T cell activation (GO:0042110), IL-4 response (GO:0070670), and WNT pathway regulation via RUNX3 (R-HSA-8951430, R-HSA-3769402), consistent with methylomic enrichment. Metabolites such as LysoPC(22:5) and phosphatidylethanolamine contributed positively, whereas carnitines, phosphatidylcholine, and caffeine metabolites (5-acetylamino-6-amino-3-methyluracil, 4-methylcatechol) were negative; urinary L-glutamine reflected increased amino acid synthesis.

Factor 8, linked to interferon responses, included whole-blood negative contributors in interleukin-27 signaling (GO:0070106), NAD/NADP salvage (R-HSA-196807, R-HSA-197264), interferon-induced stress, apoptosis, and activation (R-HSA-73893, GO:0140467, GO:0006915). B-cell transcriptomes overlapped with class I MHC antigen presentation (R-HSA-983169) and hypoxia-related stress/apoptosis pathways (R-HSA-1234174, R-HSA-5357801), with WNT signaling (R-HSA-195721) negatively contributing, consistent with its protective role in factor 4. Plasma negative metabolites included N1-methyl-2-pyridone-5-carboxamide, LysoPC(22:5), undecanedioic acid, and isoleucylproline, whereas phosphatidylcholine PC(18:1(9Z)/0:0), precursor of LysoPC(22:5) and lactosylceramide, were positive contributors.

Factor 10 also reflected interferon-associated pathways, with positive contributors enriched for Interferon Signaling (R-HSA-909733), cytokine response (GO:0034097), and NAD⁺ salvage (R-HSA-197264). B-cell contributors included interferon signaling and regulation of proliferation (GO:0042127), while negative contributors highlighted ER-stress and intrinsic apoptosis (R-HSA-380994, GO:1902235). Metabolomics positive contributors included 1-hydroxyibuprofen and palmitoylcarnitine, whereas negative contributors were PC(16:0/22:5), sphingosine 1-phosphate (d19:1-P), and sphingomyelin (d18:1/14:0), consistent with active S1P metabolism as part of the lysophospholipid signaling network.

Importantly, LPA LysoPC(22:5) emerged as the top metabolite contributor to factors 4 and 8, linking interferon-driven activation to lipid signaling pathways. In parallel, S1P, a closely related lysophospholipid mediator, contributed to factor 10, suggesting coordinated regulation of lysophospholipid metabolism in SjD. Given the strong multi-omics association of LPA with immune activation, the expression of its main signaling receptor, LPAR6, was next examined specifically in B cells to assess whether this pathway could directly influence B-cell function.

### 3.6 Validation of LPAR6 expression on B lymphocytes

To validate that circulating B cells can sense LPA signals, the LPAR6 protein was measured by flow cytometry (**Figure 4A**). B cells isolated from apheresis residues were stained with anti-LPAR6 antibody together with CD19 to confirm B cell identity. Flow cytometry analysis demonstrated LPAR6 expression on CD19⁺ B lymphocytes, as evidenced by a clear shift in fluorescence compared with the corresponding isotype and secondary-only controls (**Figure 4B**). Statistical analysis by the Kruskal–Wallis rank sum test confirmed a significant difference in the percentage of CD19⁺ and LPAR6⁺ gated cells (p = 0.02651; n = 3; **Figure 4C**). These results demonstrate that circulating B cells express LPAR6 protein and are therefore potentially responsive to plasma LPA signals, further supporting a role for the LPA–LPAR6 axis in the immunopathogenesis of SjD.

**Figure 4.**
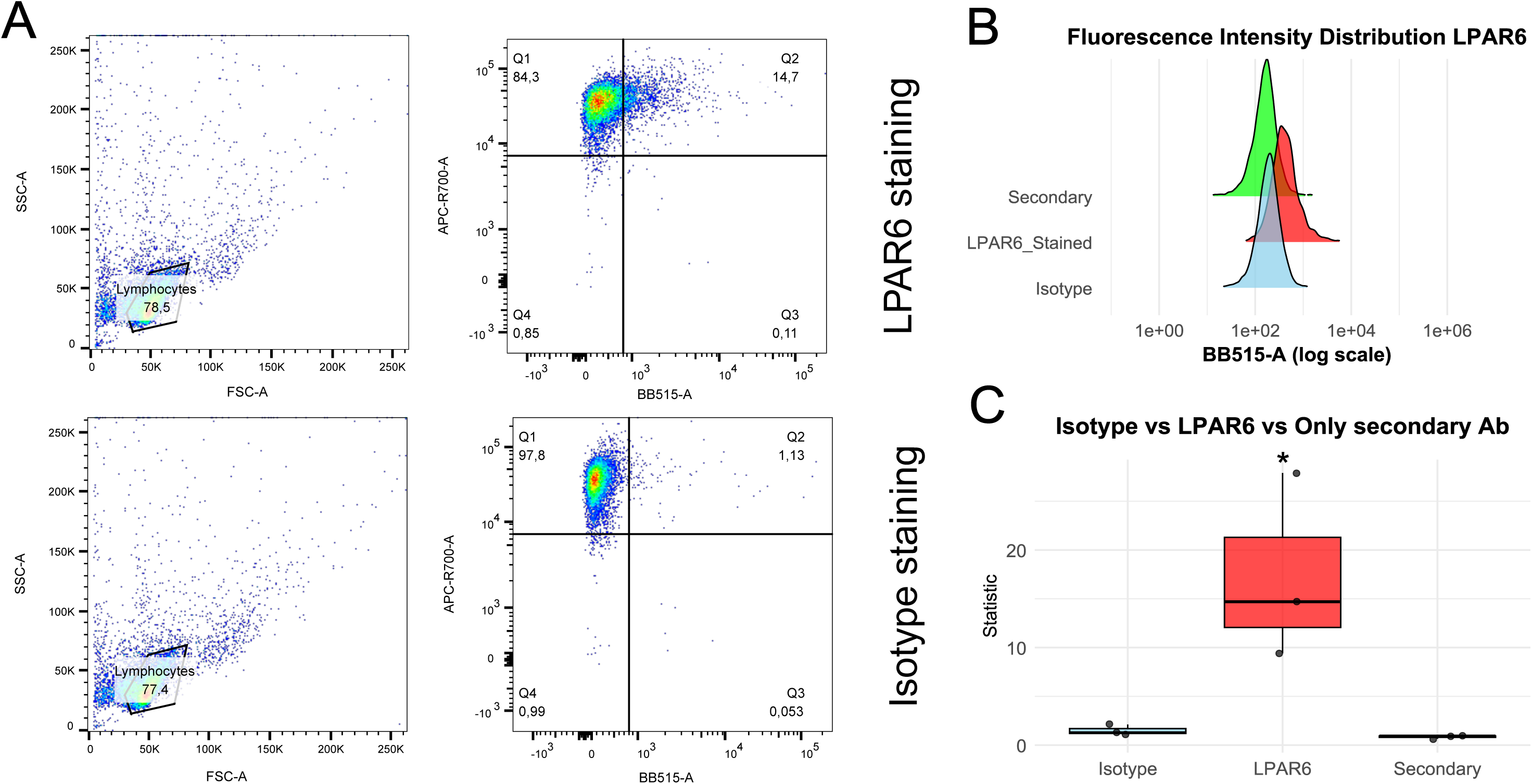
Validation of LPAR6 protein expression in B cells. **A)** Flow cytometry gating strategy used to identify B cells (APC-R700-A channel) and measure LPAR6 signal (BB515-A channel). **B)** Assessment of LPAR6 antibody specificity by comparison with isotype control, secondary antibody–only control, and LPAR6-stained samples. **C)** Boxplot showing the percentage of CD19^⁺^/LPAR6^⁺^ gated cells. Statistical comparison performed using the Kruskal–Wallis rank-sum test (*p < 0.05).

## 4. Discussion

The findings of this study are summarized in **Figure 5**.

**Figure 5.**
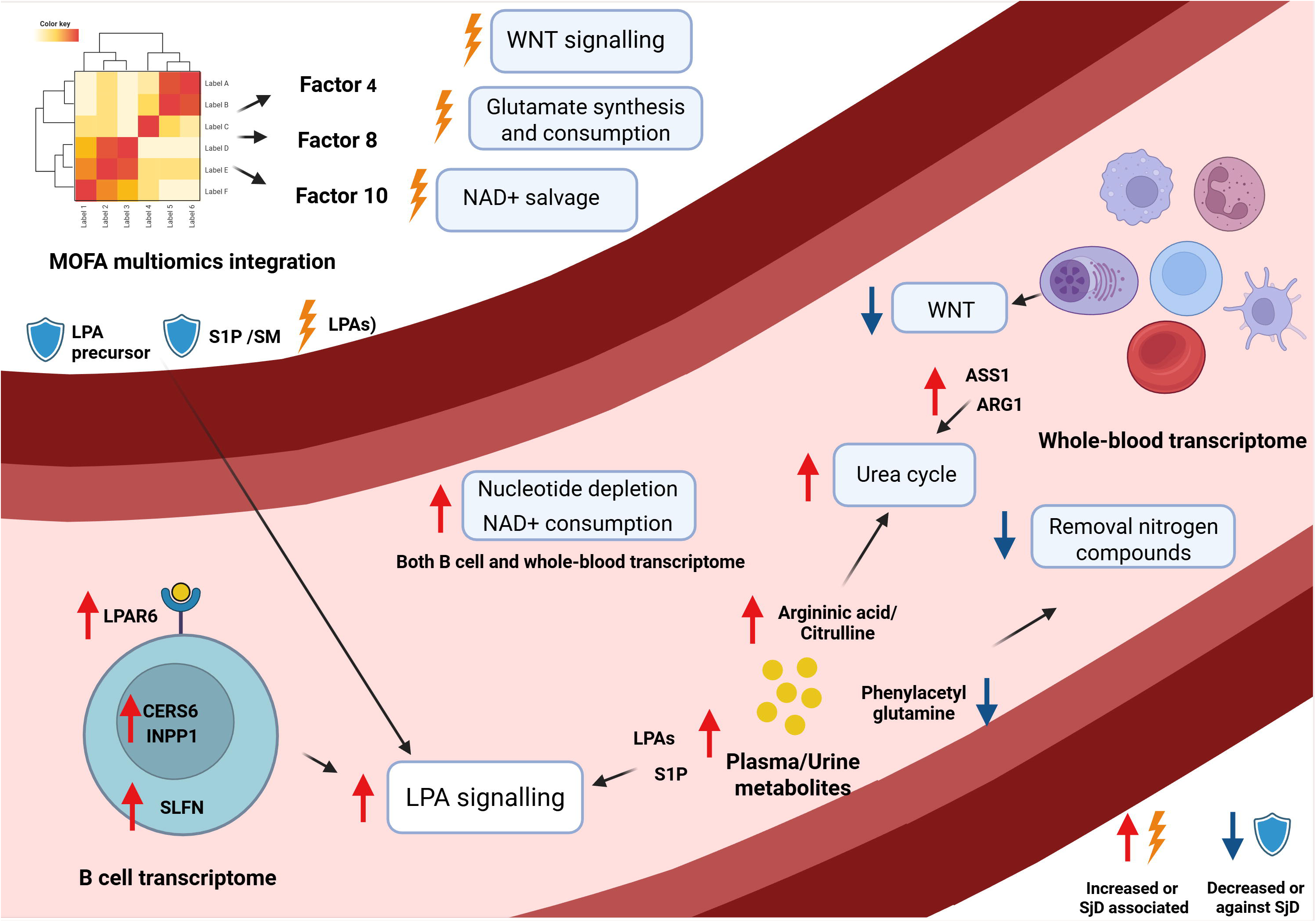
Graphical representation of intra-cellular signalling and metabolism pathways and their connection in association with metabolic changes in the macroenvironment. The figures represent the B cell and whole blood transcriptome, the plasma and urine metabolomics, and the MOFA multiomics integration results. Boxes indicate the biological pathways altered in the SjD condition, associated with an increase (red arrow) or reduction (blue arrow) of the gene expression or metabolites. From the MOFA analysis, contributors or pathways associated with the SjD phenotype (thunderbold) or against the SjD phenotype (shield) are shown.

### 4.1 The Metabolic Macroenvironment of Sjögren’s Disease: Nitrogen and Redox Stress

Our multi-omics analysis depicts a SjD macroenvironment that is metabolically rewired, nutrient-hungry, and pro-inflammatory. One of the clearest signatures involves nitrogen metabolism. Urinary phenylacetylglutamine depletion, together with elevated 2-phenylglycine, suggests diversion of nitrogen disposal pathways when urea cycle function is stressed[30]. This is complemented by increased plasma citrulline and upregulation of ASS1 and ARG1 in whole-blood transcriptomes, consistent with heightened arginine–ornithine–citrulline cycling. Together, these data indicate that SjD immune cells operate under conditions of active nitrogen turnover, likely to support proliferation and effector functions. MOFA Factor 4 reinforces this interpretation, associating cellular activation with glutamine utilization and arginine metabolism through positive contributors such as GLUL (glutamine synthetase) and ARG1. The presence of PADI2 and PADI14 among positive contributors suggests conversion of arginine to citrulline, generating citrullinated autoantigens known to trigger immune recognition in autoimmune settings[31]. Reduced creatine in patients without nephropathy may reflect muscle mass loss and SjD-related fatigue[32,33] rather than renal dysfunction[34], aligning with a systemic catabolic state.

In parallel, whole-blood transcriptomics and MOFA analysis reveal enhanced NAD⁺ metabolism and salvage pathways, indicative of increased redox stress and DNA repair demand. Genes encoding PARPs (PARP9, PARP10, PARP14) and the NADase CD38 were markedly upregulated, suggesting NAD⁺ depletion through immune activation. Factor 8 linked NAD⁺ metabolism to interferon-driven inflammation, with negative contributors in both whole blood and B lymphocytes, including NAD⁺ salvage enzymes and the NAD⁺ breakdown product N1-methyl-2-pyridone-5-carboxamide detected in plasma. Factor 4 further implicates NAD+ metabolism, with NAMPT, a salvage pathway rate-limiting enzyme, and BST1, linked to the Preiss-Handler pathway via microbiota, as positive contributors[35]. These findings together depict a macroenvironment under sustained metabolic pressure, depleting amino acids, nucleotides, and NAD⁺ to sustain immune activation.

### 4.2 Immune Activation and Interferon Signatures

This metabolically primed environment is paralleled by transcriptional evidence of immune activation. Whole-blood transcriptomes showed robust upregulation of interferon-stimulated genes, activation of Toll-like receptor signaling, macrophage activation, antigen presentation, and apoptosis pathways. GSEA confirmed induction of type I and type II interferon pathways as well as signatures of oxidative and ER stress. The MOFA analysis highlighted the discriminant factors 8 and 10 as IFN response driven, supported by the correlation with many autoantibodies including SSA and SSB, known to induce and sustain IFN response[36].

Notably, we observed overexpression of RSAD2 and CMPK2, enzymes that deplete CTP/UTP by converting them to ddhCTP[37], thereby inhibiting pyrimidine biosynthesis and triggering antiviral-like inflammatory responses[38]. Together with TYMS, NT5C3A, TYMP, TK1, and ASS1, these findings reveal a state of pyrimidine depletion without compensatory nucleotide rescue, a contrast to SLE B cells, where such rescue pathways are activated. This suggests that in SjD, B cells may remain in a sustained stress-prone state, amplifying interferon and inflammatory signaling rather than resolving it. Many similarities also arise in the MOFA factor calculated and metabolic pathways activated between SjD and SLE, allowing for comparison of similarities and differences among them (**Figure 6**).

**Figure 6.**
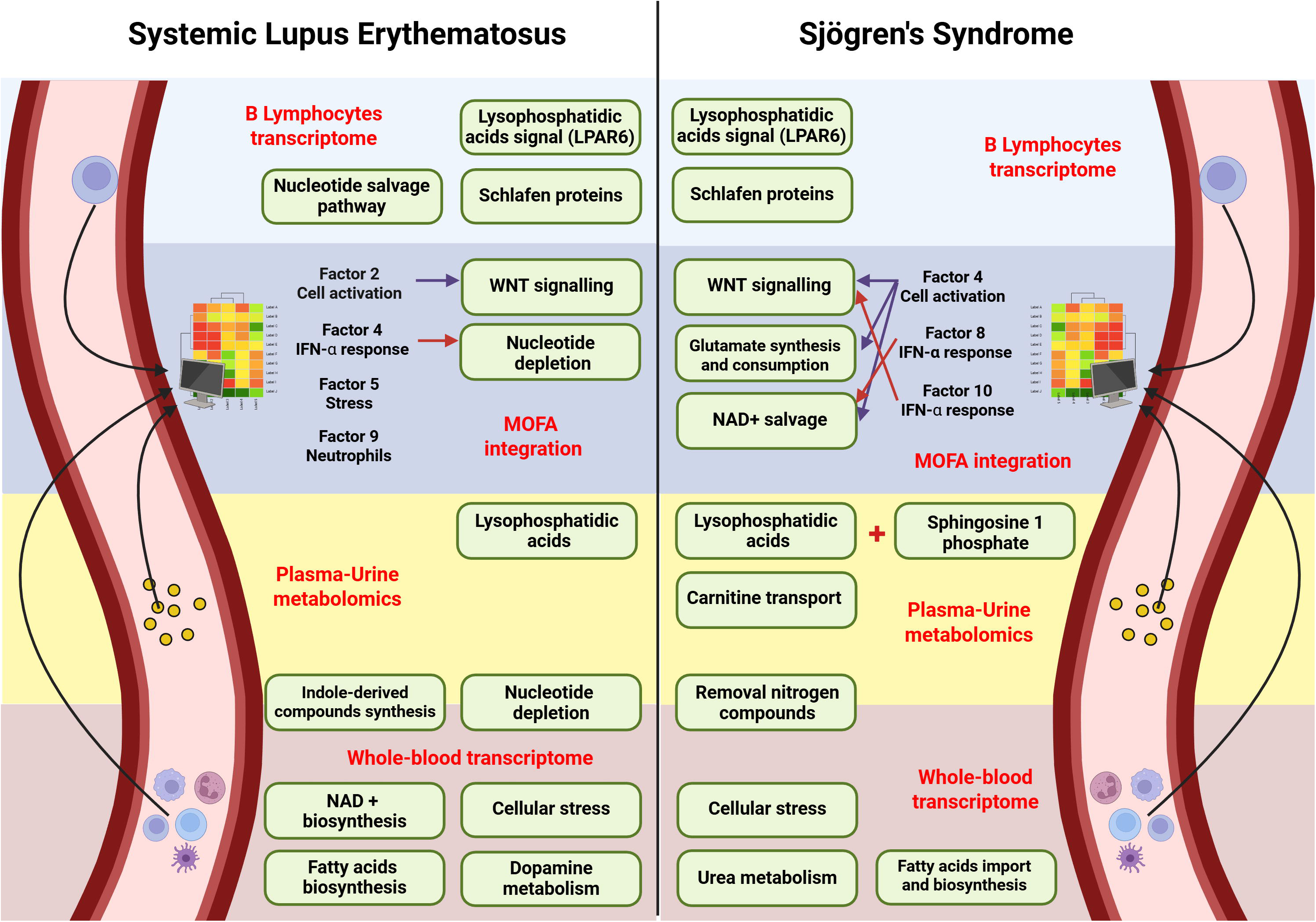
Overview of increased intracellular signaling and metabolic pathways identified in SLE and SJS. This comprehensive analysis includes data from the whole blood and B lymphocyte transcriptomes, plasma and urine metabolomics, and multi-omics integration factors calculated by the MOFA algorithm. Each dataset is distinctly color-coded in the background. Metabolic and signaling pathways are highlighted within a green box, positioned according to the dataset from which the observations are derived.

#### LPA and S1P as Lipid Messengers Linking Metabolism to Immunity

Within this nitrogen- and redox-stressed immune environment, our multi-omics approach identified bioactive lipids as key mediators connecting metabolism to immune activation. Plasma metabolomics revealed increases in LPA and S1P, both potent immunomodulatory lipids[39–41]. LPA is produced by autotaxin (ENPP2) and signals through six G-protein–coupled receptors (LPAR1–6), while S1P, its metabolic cousin, is derived from sphingomyelin and signals through S1PR1–6. These lipids trigger diverse protein activations to modulate immune responses. For instance, ENPP2 knockout models show reduced CD8+ T cell cytotoxicity[28]. LPAs also stimulate IL-10 production in dendritic cells, suggesting anti-inflammatory effects[42]. Similarly, S1Ps regulate immune responses through mechanisms like IDO1 induction and Treg differentiation in cocultures of human amniotic stem cells and PBMCs[43]. Both LPA and S1P are targets of autoimmune therapies. As previously mentioned LPA1 receptor inhibitor is in phase III clinical trials for scleroderma while S1P receptor modulators are already approved for multiple sclerosis (e.g., fingolimod) acting on S1P receptor internalisation and degradation to prevent T cells migration in the central nervous system[44]. These developments highlight their prominent roles in autoimmune diseases and suggest they may also be relevant in SjD. This is further supported by the known S1P active role in the marginal zone B cell migration[45], a process fundamental in SjD pathogenesis [45].

MOFA Factor 8 placed LysoPC(22:5), a major LPA precursor, as the top contributor associated with interferon signaling, positioning LPA signaling as contributor or effector of inflammatory response. Factor 4 also captured LysoPC(22:5) as a positive contributor, linking it to cellular activation, whereas S1P and its precursor sphingomyelin contributed negatively to Factor 10, suggesting a potential counter-regulatory effect. Transcriptomic analysis of B cells provided complementary evidence, showing upregulation of LPAR6, TRIP6 (an LPA-dependent integrin adaptor), APOL6, and CERS6, collectively indicating that SjD B cells are poised to sense and respond to LPA and sphingolipid signals. This is further supported by the overexpression of LPAR6 in SjD B lymphocytes, mirroring findings in SLE[13]. (**Figure 6**). Given the centrality of this finding, the LPAR6 protein expression on circulating B cells was experimentally validated, confirming that they indeed express this receptor and are likely responsive to LPA signals in vivo.

Together, these results suggest that the LPA–LPAR6 axis represents a key metabolic–immune bridge that may amplify a vicious loop of amplification signalling involving interferon responses and perpetuating B-cell activation in SjD.

### 4.5 Additional Mechanistic Insights

Beyond the central metabolic and lipid pathways, SjD B cells exhibit additional molecular alterations that likely modulate disease activity. MOFA analysis highlighted carnitines as having dual roles: shorter-chain acylcarnitines (acetyl- or propionyl-) contributed against SjD phenotype in Factor 4, whereas long-chain species (e.g., palmitoylcarnitine) were associated with pro-pathogenic effects in Factor 10. This aligns with previous PRECISESADS findings, which implicated carnitine metabolism in SjD immune modulation [19]. Genes involved in tryptophan degradation were also identified, suggesting enhanced catabolism along this pathway[46].

B-cell transcriptomics revealed upregulation of Schlafen proteins (SLFN5, SLFN11, SLFN12L, SLFN13), paralleling observations in SLE B cells[13], and pointing to selective roles in interferon-driven transcriptional remodeling. Finally, MOFA highlighted the WNT signaling pathway as a modulatory factor: impairment of WNT appeared to oppose the SjD phenotype in Factor 4 but contributed in Factor 8. Similar patterns have been observed in rheumatoid arthritis and ankylosing spondylitis, where WNT inhibition improves disease outcomes[47]. Among the SjD phenotype opposing actors, RUNX3, a known WNT inhibitor, emerged as a negative contributor in Factor 4. Methylation of RUNX3, conversely a positive contributor, was associated with Factor 4, suggesting that epigenetic regulation limits WNT inhibition and shapes disease-relevant signaling[48]. Collectively, these additional pathways, carnitine metabolism, tryptophan degradation, Schlafen proteins, and WNT signaling add nuanced layers of regulation that complement the metabolic, interferon, and lipid-mediated mechanisms shaping the SjD B-cell environment.

## 5. Conclusions

Taken together, multi-omics analyses depict SjD as a disease of metabolically and functionally rewired B cells and the environment. Metabolomics reveal heightened nitrogen use, altered urea-cycle enzymes (ASS1, ARG1), glutamine turnover, pyrimidine depletion (RSAD2, CMPK2, TYMS, TK1), and NAD⁺ salvage/PARP-CD38 upregulation linking metabolic stress to interferon activation. Lysophospholipids, particularly plasma LPA LysoPC(22:5/22:6) and S1P, and upregulated B-cell LPAR6 and LPA-responsive genes (TRIP6, APOL6, CERS6), suggest coordinated signaling. Limitations include the lack of direct mechanistic validation of LPA–LPAR6 in B cells and the need for functional confirmation of S1P’s role. Furthermore, despite the LPAR6 protein expression was confirmed in human B cells, a comparison between CTRLs and SjD is missing. Nonetheless, these findings highlight LPA–LPAR6 as a mechanistic link to interferon-driven B-cell activation, offering a potential target to modulate autoimmunity paralleling strategies explored in cancer and fibrotic diseases.

## Acknowledgments

The research leading to these results has received support from the Innovative Medicines Initiative Joint Undertaking under Grant Agreement Number 115565, resources of which are composed of financial contributions from the European Union’s Seventh Framework Program (FP7/2007–2013) and EFPIA companies in-kind contributions. We would like to acknowledge 3TR and the PRECISESADS consortium, which guaranteed the availability of the data used in this article and the opportunity to analyse them. CI was funded by the Université de Brest and the Région Bretagne. A.F.O. acknowledges the grant RYC2024-050119-I funded by MICIU/AEI/10.13039/501100011033 and by the FSE+.

## Data and code availability

All data included in our study were analysed in R and are available upon request at ELIXIR Luxemburg, with the permanent link: doi:10.17881/th9v-xt85. Code is available at https://github.com/IxI-97/SJS_PRECISESADS.

## Declaration of generative AI and AI-assisted technologies in the manuscript preparation process

During the preparation of this work the author(s) used ChatGpt to proofread the manuscript. The authors reviewed and edited the content as needed and take full responsibility for the content of the published article.

## Notes

### Competing Interest Statement

The authors have declared no competing interest.

