## Supplementary_material for "BiomiX-Driven Multi-Omics Integration of PRECISESADS Data Reveals Lysophosphatidic Acid and Metabolic Pathway Signatures in B Cells and Immune Macroenvironment in Sjögren’s Disease": Supplementary_tables_figures.docx

**Supplementary data**

**Table S1**. B-cell samples from Sjögren’s disease and controls of the PRECISESADS cohorts were excluded from further analysis on the basis of the criteria used to filter the samples.

| **Individuals** | **Unfiltered samples** | **B cells with digital purity <90 % in MCP or** | **IFN-α-positive CTRL samples** | **Total filtered samples** |
| --- | --- | --- | --- | --- |
| **CTRL** | 27 samples | 4 samples | 4 samples | 19 samples |
| **Total SjD** | 41 samples | 13 samples |  | 28 samples |

CTRL: controls; MCP: Microenvironment Cell Populations; SjD: Sjögren’s disease

| Sample | Peak | Annotation | m/z | Ppm error | MOFA factor |
| --- | --- | --- | --- | --- | --- |
| Plasma | peak 257 | Citrulline levels | 193.1311 | 9 | No detected |
| Plasma | peak 346 | Sphingosine−1−phosphate | 380.2522 | 10 | No detected |
| Plasma | peak 412 | LysoPC(22:6) | 606.3019 | 11 | No detected |
| Plasma | ‍peak 90 | LysoPC(22:5) | 592.3319 | 1 | 1° negative in contributor Factor 8  8° positive in contributor Factor 4 |
| Plasma | peak 302‍ | Cholesterol glucuronide | 282.2019 | 4 | No detected |
| Plasma | ‍peak 323 | Octanoyl−L−carnitine | 310.1997 | 3 | No detected |
| Plasma | ‍peak 288 | Hexanoyl−L−carnitine | 260.1856 | 0 | No detected |
| Plasma | ‍peak 246 | Trimethyl−Lysine | 171.1489 | 2 | No detected |
| Urine | ‍peak 45 | Methylxanthine | 167.0565 | 1 | No detected |
| Urine | peak 155‍ | Phenylacetylglutamine | 265.1193 | 0.4437 (MS/MS) | No detected |
| Urine | peak 813‍ | Sphingosine | 322.2719 | 1 | No detected |
| Urine | peak 499‍ | Trimethyl−Lysine | 189.1591 | 3 | No detected |

**Table S2**. Plasma and Urine peaks annotation. The peaks presented were annotated by m/z (MS1 annotation, if not specified elsewhere (MS/MS).

**Table S3.** Number of samples available in the study.

| **Samples** | **B-cell transcriptome** | **Whole-blood transcriptome** | **Plasma metabolome** | **Urine metabolome** | **Whole-blood methylome** |
| --- | --- | --- | --- | --- | --- |
| **SjD/CTRL** | 41/27 | 293/508 | 45/54 | 45/54 | 282/364 |
| **SjD/CTRL integrated with MOFA** | 20/27 | 41/32 | 36/45 | 36/45 | 35/37 |

The number of Sjögren’s disease (SjD) and control (CTRL) samples from the PRECISESADS cohorts is indicated, respectively, on the left and on the right. The number of samples shared between the transcriptomic and metabolomic analyses in the MOFA integration analysis is also indicated.

|  | BLymphocytes  (RNAseq) | Plasma (Metabolomic) | Urine (Metabolomic) | Whole_blood (Methylomic) | Whole_ blood (RNAseq) |
| --- | --- | --- | --- | --- | --- |
| Factor1 | 49.95 % | 0.03 % | 0.03 % | 0.63 % | 0 % |
| Factor2 | 0.05 % | 0.74 % | 0.04 % | 30.33 % | 0 % |
| Factor3 | 0 % | 0.16 % | 0.01 % | 0.01 % | 24.33 % |
| Factor4 | 0 % | 0.26 % | 0.03 % | 2.23 % | 13.89 % |
| Factor5 | 14.71 % | 0 % | 0 % | 0 % | 0.27 % |
| Factor6 | 7.67 % | 0.01 % | 0.01 % | 0.58 % | 0 % |
| Factor7 | 0.01 % | 1.03 % | 0.01 % | 0.01 % | 7.21 % |
| Factor8 | 0.43 % | 0.17 % | 0.01 % | 0 % | 6.72 % |
| Factor9 | 0.03 % | 0.01 % | 0.01 % | 7.11 % | 0.01 % |
| Factor10 | 1.93 % | 0.01 % | 0.01 % | 0 % | 3.08 % |
| Factor11 | 3.29 % | 0.06 % | 0.04 % | 0.26 % | 0.11 % |
| Factor12 | 0 % | 0.11 % | 0.34 % | 2.9 % | 0.16 % |
| Factor13 | 2.02 % | 0.14 % | 0.47 % | 0.01 % | 0.55 % |
| Factor14 | 1.78 % | 0.62 % | 0.01 % | 0.69 % | 0 % |
| Factor15 | 0.19 % | 0.01 % | 0.06 % | 2.35 % | 0 % |
| Factor16 | 0.41 % | 0.05 % | 0.01 % | 1.78 % | 0.04 % |
| Factor17 | 0 % | 0 % | 0.01 % | 0.16 % | 1.91 % |
| Factor18 | 0 % | 0.02 % | 0.18 % | 0.36 % | 1.52 % |
| Factor19 | 1.05 % | 0.02 % | 0.01 % | 0.01 % | 0.45 % |
| Factor20 | 0.2 % | 0 % | 0.01 % | 0.03 % | 1.27 % |
| **total** | 83.72 % | 3.45 % | 1.3 % | 49.45 % | 61.52 % |

**Table S4**. Percentage (%) of the variance explained values per factors among the omics in the 20-factor model. The total percentage of variance explained per omics is reported in the last row of the table.

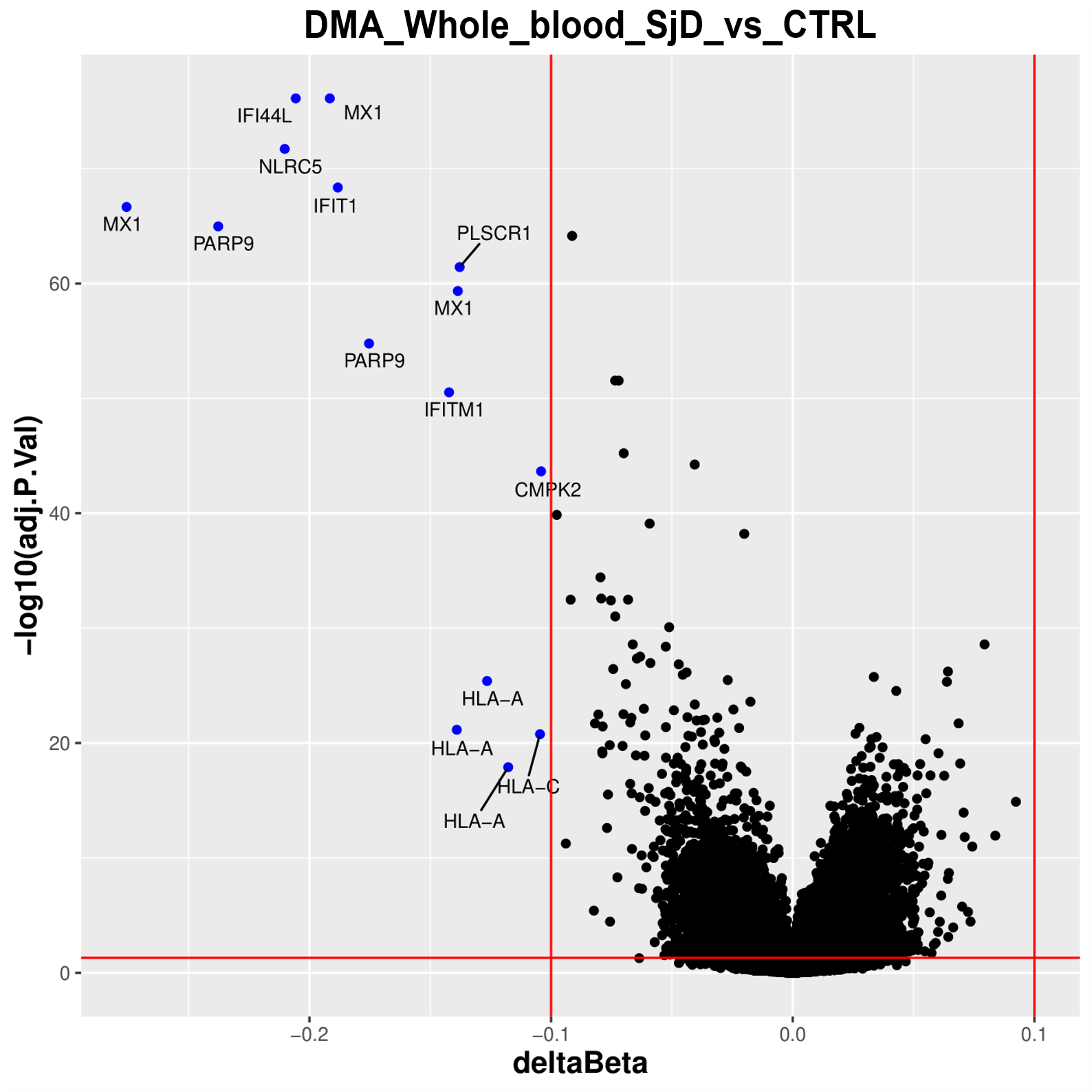
**Figure S1. Methylomics alterations in whole blood and B cells of SjD patients compared with controls (CTRL) in the PRECISESADS dataset**. **A**) Volcano plot of differential methylation analysis in whole blood. Downregulated and upregulated CpG are shown in blue. The 15 demethylated CpG genes are highlighted. Red lines indicate the thresholds of significance: |deltaBeta| > 0.1 and adjusted *p* < 0.05
